# An integrated transcriptomics and metabolomics analysis of the *Cucurbita pepo* nectary implicates key modules of primary metabolism involved in nectar synthesis and secretion

**DOI:** 10.1101/491597

**Authors:** Erik M. Solhaug, Rahul Roy, Elizabeth C. Chatt, Peter M. Klinkenberg, Nur-Aziatull Mohd-Fadzil, Marshall Hampton, Basil J. Nikolau, Clay J. Carter

## Abstract

Nectar is the main reward that flowers offer to pollinators to entice repeated visitation. *Cucurbita pepo* (squash) is an excellent model for studying nectar biology, as it has large nectaries that produce large volumes of nectar relative to most other species. Squash is also monoecious, having both female and male flowers on the same plant, which allows comparative analyses of nectary function in one individual. Here we report the nectary transcriptomes from both female and male nectaries at four stages of floral maturation. Analysis of these transcriptomes and subsequent confirmatory experiments revealed a metabolic progression in nectaries leading from starch synthesis to starch degradation and to sucrose biosynthesis. These results are consistent with previously published models of nectar secretion and also suggest how a sucrose-rich nectar can be synthesized and secreted in the absence of active transport across the plasma membrane. Non-targeted metabolomic analyses of nectars also confidently identified 40 metabolites in both female and male nectars, with some displaying preferential accumulation in nectar of either male or female flowers. Cumulatively, this study identified gene targets for reverse genetics approaches to study nectary function, as well as previously unreported nectar metabolites that may function in plant-biotic interactions.

## 1 INTRODUCTION

Floral nectar is a sugary solution that serves as a reward for pollinators and is essential for successful reproduction in many flowering plants (Simpson and Neff, 1983; Ollerton et al., 2011). Diverse species produce nectar with different relative concentrations of sugars. Arabidopsis and other members of the Brassicaceae produce nectar that is rich in hexoses (glucose and fructose) (Davis et al., 1995; Davis et al., 1996; Davis et al., 1998), whereas tobacco and squash *(Cucurbita pepo)* produce nectar rich in sucrose (Nepi et al., 2001). Squash flowers are hemitropous (Dmitruk and Weryszko-Chmielewska, 2013) with the nectaries hidden but still accessible to pollinators. *C. pepo* is a monoecious species that produces both staminate and pistillate flowers, which both secrete nectar. Nectar secretion begins at around dawn and lasts nearly 6 hours (Nepi et al., 2001). Male and female flowers in *C. pepo* differ in the timing of nectar secretion, with male flowers increasing nectar secretion until ~3 hours post flower opening before leveling off, whereas in female flowers nectar levels continue to increase throughout the morning and until the flowers start to close (at ~6 hours post-opening) (Nepi et al., 2001). It is likely that the difference in timing has biological significance as the reproductive success of a plant depends on the sequential visitation of pollinators to male flowers first to receive pollen before visiting female flowers for pollination. Also, on average female flowers produce more nectar, and it contains higher sugar content than the nectar produced by male flowers (Nepi et al., 2001). Nectar levels in both flower types decrease drastically by 6 hr post secretion, suggesting that some resorption of nectar occurs (Nepi et al., 2001).

Nectar secretion involves a number of steps that are intricately regulated in order to maximize pollination while not wasting resources (Pleasants and Chaplin, 1983; Heil, 2011). Floral nectaries in most species are non-photosynthetic sink tissues that depend on photosynthate (e.g. sucrose) and other ‘pre-nectar’ components to be delivered via the vasculature, and these are often stored prior to secretion (Pacini and Nepi, 2007; Nepi and Stpiczynska, 2008; Heil, 2011). For example, high levels of starch accumulation in the parenchyma of immature nectaries (before secretion) has been reported for many flowering species (Peng et al., 2004; Ren et al., 2007; Ren et al., 2007; Lin et al., 2014). This starch is absent from nectary tissues during and after secretion (Zhu et al., 1997; Peng et al., 2004; Ren et al., 2007; Ren et al., 2007), suggesting nectary starch may serve as a temporary carbon store to facilitate fast production of soluble sugars for nectar secretion. While starch accumulation and degradation are strongly correlated to nectar secretion in diverse plant species, the specific genes, proteins, and metabolites that are involved in this process have had limited study and only in *Nicotiana* spp. (Ren et al., 2007; Ren et al., 2007).

After starch degradation, there is a well-supported model for nectar synthesis and secretion in Arabidopsis (Lin et al., 2014; Roy et al., 2017). Specifically, starch breakdown products (chiefly maltose and hexose-phosphates) are first assembled into sucrose by the action of sucrose-phosphate synthases (SPS) and sucrose-phosphate phosphatases, among other enzymes, whereupon the sucrose is exported from the nectary cells in a concentration-dependent manner via the uniporter SWEET9. In some species that generate hexose-rich nectars, the exported sucrose may be hydrolyzed into glucose and fructose by cell wall invertases (CWINV). Evidence for this model of nectar secretion is based on biochemical analyses (Ren et al., 2007; Ren et al., 2007; Liu and Thornburg, 2012), in combination with the fact that genetic ablation of sucrose synthesis, export or extracellular hydrolysis all impair nectar secretion in Arabidopsis and/or tobacco (Ruhlmann et al., 2010; Lin et al., 2014).

Additionally, several aspects of hormonal and transcriptional control of nectary functions have been studied in Arabidopsis and other species (Heil et al., 2001; Carter and Thornburg, 2003; Liu et al., 2009; Radhika et al., 2010; Liu and Thornburg, 2012; Wang et al., 2014). However, a majority of studies, particularly in Arabidopsis, have largely been dependent on genetic strategies, and biochemical confirmation of the conclusions have been hampered by the small size of Arabidopsis flowers and small amounts of nectar produced by these flowers. In contrast, plants in the *Cucurbita* genus, such as *C. pepo,* develop much larger flowers and produce >100-fold more nectar per flower than Arabidopsis, and thus are more amenable to biochemical studies on nectar production [e.g. (Nepi et al., 1996; Nepi et al., 2001; Nepi et al., 2012; Chatt et al., 2018)].

Here we expand the potential utility of *C. pepo* and related cucurbits as models to pursue physiological, genetic, and biochemical studies of nectar production through transcriptomic and metabolomics analyses of nectaries and nectar through the process of maturation. Subsequent experiments placed genes, pathways and metabolites in a physiological context. This report represents an important step in improving our understanding of cucurbit nectary biology.

## 2 METHODS

### 2.1 Plant growth, tissue collection, RNA isolation and sequencing

*Cucurbita pepo* (Crookneck Yellow Squash) plants were grown on Sun Gro LC8 soil under a 16 hr day/8 hr night cycle, photosynthetic photon flux of 250 μmol m^-2^ s^-1^ at leaf level, and a temperature of ~23°C. Four types of RNA samples were separately prepared from manually collected nectaries of both male and female squash flowers, including: ‘pre-secretory #1’ (24 hours prior to anthesis/nectar secretion), ‘pre-secretory #2’ (15 hours prior to anthesis/nectar secretion), ‘secretory’ (full anthesis, 2-3 hours after dawn), and ‘post-secretory’ (9 hours after the ‘secretory’ stage). All nectary tissues were manually dissected by hand with the RNA being immediately extracted by mechanical disruption with a microcentrifuge pestle and using an RNAqueous^®^ RNA isolation kit (Ambion, Austin, TX) with Plant RNA Isolation Aid (Ambion, Austin, TX). Agarose gel electrophoresis and UV spectrophotometry were used to assess RNA quality for all samples prior to submission to the University of Minnesota Genomics Center for mRNA isolation, barcoded library creation and Illumina HiSeq 2500 sequencing. Twenty-three TruSeq RNA v2 libraries were created (triplicate samples for male and female nectaries at four timepoints each, except for only duplicate samples of female ‘pre-secretory #2’ nectaries) and sequenced via 50 bp, paired-end runs on the HiSeq 2500 using Rapid chemistry. All libraries were pooled and sequenced across two full lanes. This generated over 240 M reads for each lane and the average quality scores were above Q30.

### 2.2 Informatic analyses

The sequenced reads from nectary samples were assembled separately using Trinity (Grabherr et al., 2011), which automatically takes read quality and consistency into account, and yielded 17,772 contigs. This contig set was mapped to both the *Cucumis melo* and *Arabidopsis thaliana* genomes by NCBI’s blastn, with an E-value cutoff of 0.00001, and the counts upper-quartile normalized. Normalized counts were fitted to a negative binomial distribution using DESeq v1.6.1 (Anders and Huber, 2010). The resulting p-values from DESeq were filtered by restricting to contigs with a 50% or greater change in mean expression between nectary stages. The Benjamini-Hochberg method was used to control the false discovery rate of contigs determined to be differentially expressed to 0.05 (Benjamini and Hochberg, 1995).

For gene ontology analyses, genes displaying >2-fold and statistically significant differences (as determined by DESeq and FDR via the Benjamini-Hochberg method described above, p < 0.05) in various stages of only female, only male or both nectaries combined were analyzed using the ‘Statistical Overrepresentation Test’ tool via the PANTHER Classification System using the default settings and Bonferroni correction for multiple testing (p < 0.05) [(Mi et al., 2017) http://pantherdb.org/]. Information on biological processes, molecular function and cellular component terms were extracted and saved in Microsoft Excel sheets. Heatmaps representing gene expression of candidate genes in male and female nectaries was performed using the Heml 1.0 (Heatmap Illustrator) software package. Fold change for each candidate gene for each stage was calculated with respect to the 0 hr stage. The ratios were then used by the Heml toolkit to generate heatmaps. Scales were kept the same for all genes analyzed across various stages of both male and female flowers

### 2.3 Data availability

Raw sequence reads are available at the National Center for Biotechnology Information Sequence Read Archive under GEO accession number GSE111695.

### 2.4 Real time RT-PCR (RNAseq validation)

The same RNA samples used for sequencing were also subjected to cDNA preparation using the Promega GoScript Reverse Transcription System (Catalog # A5000), with 1 μg of RNA used for cDNA preparation. Expression patterns for key genes that showed differential expression via RNAseq analyses were validated by quantitative RT-PCR using Agilent Brilliant III Ultra-fast SYBR Green QPCR Master Mix (Catalog #600882) and a cDNA template concentration of 1 ng/μl. Expression values are expressed as fold change relative to 0 h timepoint and are based on the delta delta Ct values obtained from the normalized Ct values for each gene. Gene expression was normalized to a gene encoding a RING/U-Box ligase superfamily protein (C. melo hit= gi|65907532 8|ref|XM _008439865.1|). This gene was chosen as the internal reference based on its stable expression level in all nectary samples in our RNA-seq dataset. Primer sequences for each gene are provided in Supplemental Table 1).

### 2.5 Lugol staining

Male and female squash flowers were bisected longitudinally with a scalpel. The flower halves were dipped in 0.05 Molar Iodine/Potassium iodide stain (Fischer Scientific Cat#S93408) for 60 seconds. They were subsequently washed twice for 2 minutes each in 50 ml of water to remove excess stain and imaged using a dissecting microscope. Care was taken not to have an air bubble trapped in the nectary cup as the flower was dipped in the lugol solution.

### 2.6 Quantitative starch assay

Total starch was quantitatively determined using the Megazyme Total Starch Assay Kit Analysis (Megazyme, Cat# K-TSTA). Samples were ground in 0.25 ml of 80% ethanol, then an additional 0.25 ml of 80% ethanol was added. Samples were incubated at 80°C for 5 min in order to remove soluble sugars and maltodextrins. Samples were centrifuged for 10 min at 4300 rpm and supernatants were removed. The samples were resuspended in 1 ml 80% ethanol followed by repeating centrifugation at 4300 rpm for 10 min and discarding supernatant. 300μl of α-amylase (100 U/ml in 100 mM sodium acetate buffer, pH 5.0) was added to each sample, followed by incubation at 100°C for 6 min with vortexing at 2 min intervals. 10 μl of amyloglucosidase (3,300 U/mL) was then added to each sample and samples were incubated at 50°C for 30 min. Samples were then made up to 1 mL final volume then centrifuged at 3000 rpm. Supernatants were then assayed for D-glucose using the Glucose oxidase/peroxidase (GOPOD) method. The – 24 hr and -15 hr samples were diluted 10-fold in order to get the concentration of D-glucose in the range of detection for the GOPOD assay. 33 μl of supernatant was added to 1 ml of GOPOD solution (made according to kit protocol) in duplicate and tubes were incubated at 50°C for 20 min. Absorbance was read at 510 nm using a visible spectrophotometer. Starch content (% w/w) was calculated using the equation for solid samples provided in the megazyme protocol supplied with the kit.

### 2.7 Beta-amylase (BAM) activity assays

Nectary tissue was harvested at all four stages of nectar secretion and stored at -80°C before use. Samples were ground in 150 μl Tris-HCl, pH 8.0 on ice then centrifuged at max speed for 20 minutes at 4°C. 10 μl of supernatant was added to a new tube containing 150 μl of 0.1 M sodium acetate (pH 4.6) and 75 μl of amylopectin (20 mg/ml in 0.2 M KOH). Each reaction was then made up to 300 μl with deionized water and then incubated at 37°C for 60 min. The reactions were stopped by incubation at 100°C for 3 min. 20 μl of each reaction was added to 750 μl of PAHBAH solution and DI water was added to a final volume of 1 ml. Samples were then incubated at 100°C for 5 min then absorbance at 410 nm was read for each sample blanked against a reagent blank with Tris-HCl grinding buffer added in place of crude protein.

Amylopectin stocks were prepared by boiling Amylopectin/KOH solution for 10 min until the solution became clear, then aliquots were stored at -20°C prior to use. PAHBAH solution was prepared by mixing 1 part 5% p-hydroxybenzoic acid hydrazide in 0.5 M HCl with 4 parts 0.5 M NaOH.

### 2.8 Soluble sugar assay

Soluble sugars were quantified using an Amp-red/glucose oxidase/horseradish peroxidase method coupled with endogenous invertase activity as previously described (Ruhlmann et al., 2010; Bender et al., 2012). In brief, nectary tissue from each stage was ground in protein extraction buffer then subjected to 20 min treatment with and without addition of 10 mM sucrose. Reactions were diluted 1:100 to get into the working range of the glucose assay. The glucose assay mix (Ruhlmann et al., 2010) consisted of 38.5 μM Amp-red (stock solution dissolved in DMSO), 5 units of horse radish peroxidase and glucose oxidase, and 33 mM sodium phosphate buffer pH 7. 25 μl of assay solution was mixed with 75 μl of diluted sugar solution and absorbance was measured at 570 nm for each sample. Values were compared to a standard curve with glucose to determine the absolute amount of soluble sugar in each sample.

### 2.9 Nectar metabolomics

Two separate GC-MS based methods were employed for untargeted metabolite profiling of six nectar samples from independent male and female flowers of *C. pepo.* The first of these provided data on the predominant sugars that constitute the nectar (i.e. sucrose, glucose, and fructose). Specifically, 1 μL of nectar was spiked with 10 μg ribitol as an internal standard, and the mixture was dried overnight by lyophilization. The sample underwent methoximation at 30 °C for 90 min while continuously shaking with 20 mg mL^-1^ methoxyamine hydrochloride dissolved in pyridine. The methoximated sample was silylated for 30 min at 60 °C with BSTFA/ 1% TCMS. Following dilution with 1.5 mL pyridine, 1-μL of sample was analyzed by GC-MS.

Less abundant constituents of the nectar were extracted from a 5-μL aliquot of nectar sample that was spiked with 0.5 μg nonadecanoic acid and 1 μg ribitol as internal standards. Hot methanol (2.5 mL) was immediately added to the nectar, and the mixture was incubated at 60 °C for 10 min. Following sonication for 10 min at 4 °C, chloroform (2.5 mL) and water (1.5mL) were sequentially added, and the mixture was vortexed. Centrifugation separated the polar and non-polar fractions, and the entire non-polar fraction and half of the polar fraction was recovered to separate 2 mL screw-cap glass vials and dried overnight by lyophilization. The polar fraction underwent methoximation as previously described, and both polar and non-polar fraction were silylated for 30 min at 60 °C with BSTFA/ 1% TCMS.

The derivatized metabolites (the sugars, polar, and non-polar fractions) were analyzed using an Agilent Technologies Model 7890A GC system equipped with an HP-5ms (30 m, 0.25 mm, 0.25 μm) GC column that was coupled to an Agilent Technologies 7683B series injector and Agilent Technologies Model 5975C inert XL MSD with Triple-Axis Detector mass spectrometer. Chromatography parameters for the polar and non-polar fractions were set to a helium gas flow rate of 1 mL min^-1^, 2 μL injection, with a temperature gradient of 80 °C to 320 °C increasing at a rate of 5 °C min ^-1^, followed by a 9 min hold at 320 °C. The polar fractions were analyzed using a “heart-cut” method (Boeker et al., 2013) which diverted gas flow to an FID detector during elution times for fructose, glucose, and sucrose.

GC parameters for predominant sugar metabolites were set to a helium gas flow rate of 1 mL min^-1^, 1 μL injection with a 10:1 split, and a temperature gradient of 100 °C to 180 °C increasing at a rate of 15 °C min ^-1^, then 5 °C min ^-1^ to 305 °C, then 15 °C min ^-1^ to 320 °C, followed by a 5 min hold at 320 °C. Deconvolution and integration of resulting spectra was performed with AMDIS (Automated Mass Spectral Deconvolution and Identification System) software. Analyte peaks were identified by comparing mass spectra and retention indices to the NIST14 Mass Spectral Library and authentic standards when possible to confirm identification.

## 3 RESULTS

### 3.1 Experimental Design

To understand the nexus of genes and processes that are activated in *C. pepo* nectaries throughout maturation, we decided to take a transcriptomic strategy. This approach was previously used with Arabidopsis nectaries (Kram et al., 2009) and led to the identification of key genes that regulate nectary function and nectar production [e.g. (Kram and Carter, 2009; Ruhlmann et al., 2010; Bender et al., 2012; Bender et al., 2013; Lin et al., 2014; Wiesen et al., 2016; Schmitt et al., 2018)]. Squash nectaries are considerably larger than Arabidopsis (~1 cm diameter for squash vs. ~100 μM for Arabidopsis) and produce relatively large amounts of nectar (>50 μl for squash vs. <<1 μl for Arabidopsis, per flower). A transcriptomic understanding of the squash model would thus further the field of nectar biology, as physiological, biochemical and cellular studies in downstream experiments would be made easier than with other species. We proceeded with an experimental plan where four stages of both pistillate (female) and staminate (male) flowers were selected to study nectary gene expression as affected by maturation (Figure 1).

**Figure 1.**
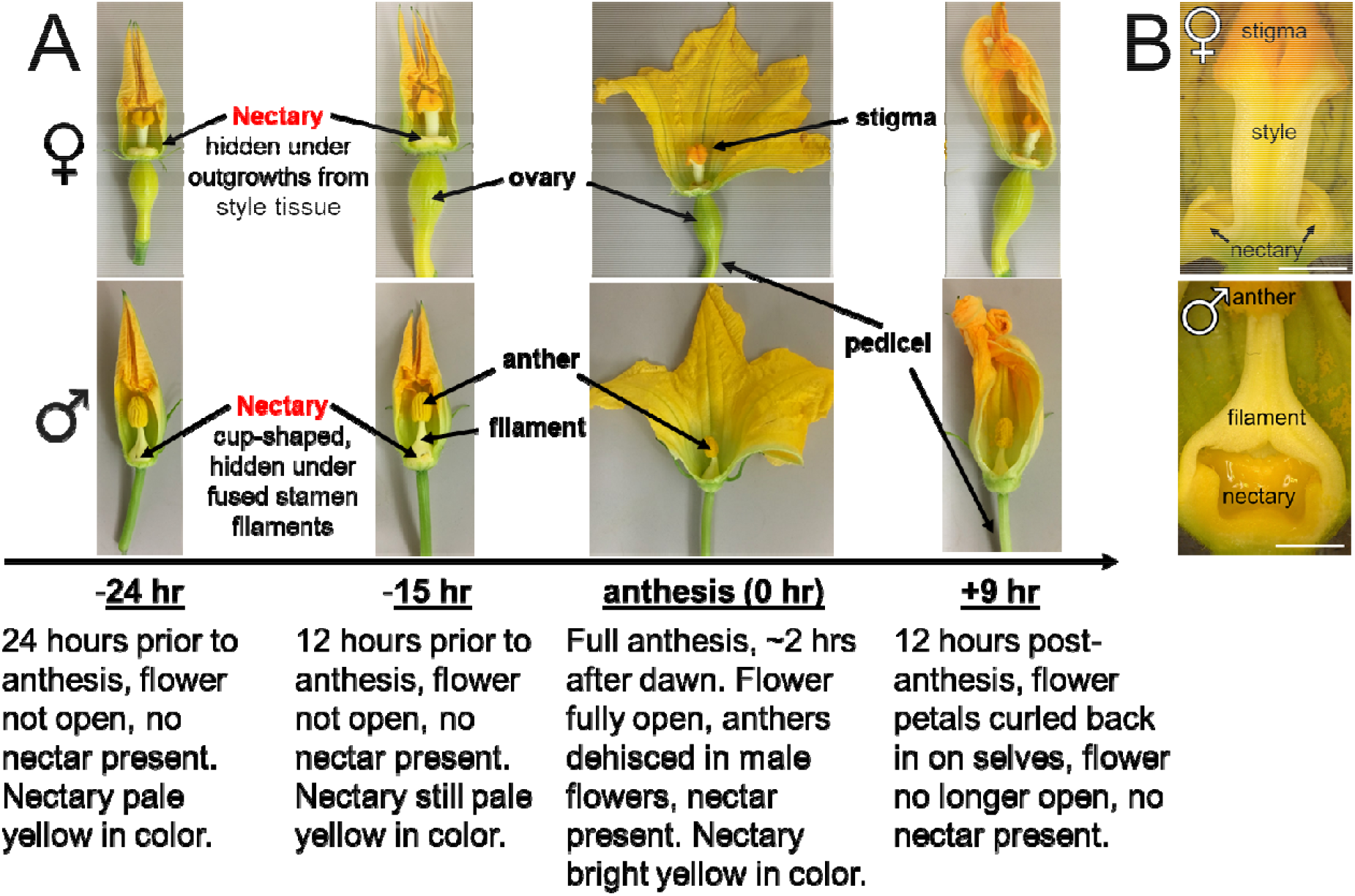
Developmental stages and tissues used for squash nectar/y analyses. (A) Time points used for female (top) and male (bottom) nectary collections. (B) Cross-section of female and male flowers to reveal nectaries. Scale bar equals 5 mm.

The developmental stages chosen for analyses centered around anthesis, which is when pollen is released by anthers. Nectar production is synchronized with anthesis (Nepi et al., 1996; Nepi et al., 2001) and hence this study was centered around the time of anthesis, which aided the uniform timing of tissue collection for subsequent studies. Squash flowers progress from small, closed green buds to open flowers relatively rapidly (within a few days), and we tracked their progress through key visual cues. The ‘pre-secretory stage 1’ occurs 24 hours before anthesis, and was recognized by the fact that buds are still closed but have a yellow coloration at the tips of the fused corolla; this stage is designated as the ‘-24 hr’ time-point (Figure 1A). We noted that nine hours later the yellow color intensifies and the tip of the corolla is slightly unfurled and we designated this as ‘pre-secretory stage 2’ or the ‘-15 hr’ time-point (~12 hours before dawn). Open flowers actively producing nectar (~3 hours after dawn) were termed ‘secretory stage’ or the ‘0 hr’ time-point, and 9 hours later (~12 hours after dawn), when the flowers had closed, was chosen as ‘post-secretory’ or ‘+9 hr’ time-point. While nectar was completely absent from the flowers at the -24 and -15 hr time-points, copious amounts of nectar were present in the 0 hr and +9 hr flowers. Replicate RNAs were isolated from the nectaries (e.g. Figure 1B) at each of these time-points and subjected to Illumina-based RNA-seq analyses.

### 3.2 RNA sequencing and differential expression analyses

Over 240M reads (50 bp, paired end) derived from squash nectary RNAs were assembled into 39,978 contigs and subsequently mapped to both the *C. melo* (melon) and *Arabidopsis thaliana* Col-0 genomes. The rationale for mapping contigs to melon is that it is a close relative to squash with a fully sequenced genome; similarly, contigs were mapped to Arabidopsis because it has a very well annotated genome and has served as the genetic model for plant biology, including the process of nectar production [reviewed in (Roy et al., 2017)]. Altogether, *C. pepo* nectary contigs mapped to 8,863 unique *C. melo* genes (Supplemental Table 1), which were subsequently used for all downstream analyses of differential expression.

Differentially expressed genes between nectary stages were defined as having a statistically significant *and* mean two-fold difference in expression. The lists of differentially expressed genes between stages in female and male nectaries are available in Supplemental File 2, respectively. We additionally performed a separate differential expression analysis where reads from both female and male nectaries were grouped together by developmental stage (Supplemental File 2). These analyses revealed genes that are upregulated at specific stages of maturation, as well as ones that were commonly expressed at all time-points in both male and female flowers (Figure 2, Supplemental Files 3-6). The numbers of genes displaying stage-dependent upregulation in nectary expression are indicated in the Venn diagrams for female (Figure 2A; n = 3 for each stage, except n = 2 for -15 hr), male (Figure 2B, n = 3 for each stage), female and male together when analyzed for significance separately (Figure 2C; i.e. differentially expressed genes commonly appearing in both Figure 2A and 2B), and female and male together when all reads for a given stage were analyzed as a group for fold-change and statistically significant differences in expression (Figure 2D; n = 6 for -24, 0 and +9 hr, n = 5 for -15 hr). The non-overlapping regions of each Venn diagram indicate the number of genes that are upregulated at a specific developmental stage (i.e., at the -24 hr, -15 hr, 0 hr or +9 hr time-points), whereas the numbers shown in overlapping regions are commonly upregulated at two or more time-points over the others. For example, in female nectaries (Figure 2A) there were 394 genes expressed >2-fold higher (and statistically significant) at both the -24 and -15 hr time-points over the 0 and +9 hr time-points; similarly, there were 209 genes that were more highly expressed at the -24, – 15 and 0 hr time-points over the +9 hr stage.

**Figure 2.**
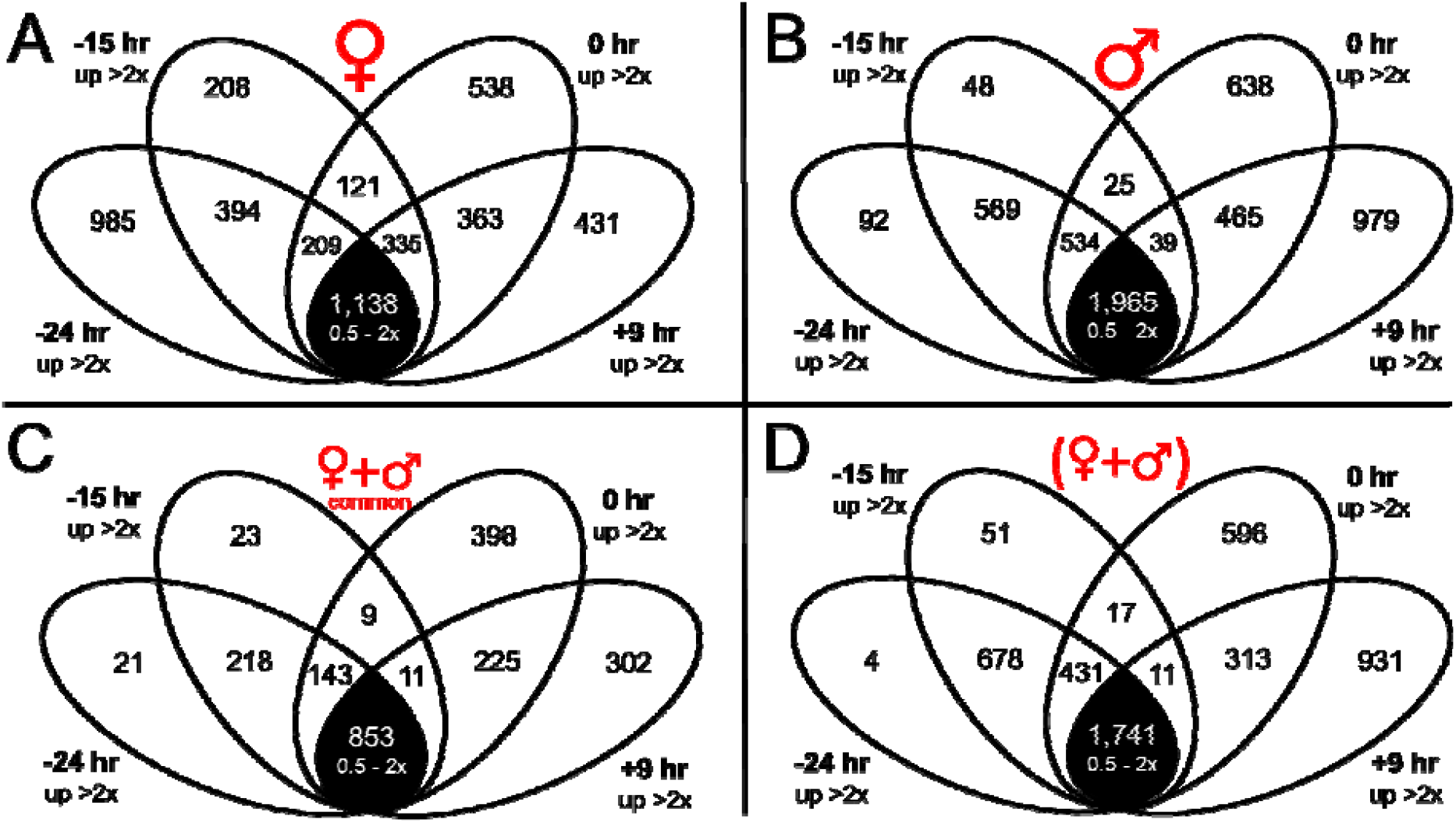
Venn diagram set-representation of differentially expressed genes in nectaries associated with developmental stages. Differentially expressed genes are defined as those showing a >2-fold change in expression in any pair-wise or multi-stage comparison (p<0.05). The number of genes that fail this test (commonly expressed at all time-points) are identified in the black-shaded subset. (A) Number of genes up-regulated in female nectaries at the -24 hr,-15 hr, 0 hr or +9 hr time-points in female nectaries. (B) Number of genes up-regulated in male nectaries at the -24 hr,-15 hr, 0 hr or +9 hr time-points. (C) Number of genes commonly upregulated in both male and female nectaries (i.e., differentially expressed genes appearing at the same time-points in both A and B) at the -24 hr,-15 hr, 0 hr or +9 hr time-points. (D) Number of genes upregulated at each developmental stage when reads from both female and male nectaries from a particular stage are analyzed together as a single unit.

### 3.3 Validation of RNA-seq data

The differential expression identified through the RNA-seq analyses was subsequently validated by qRT-PCR using RNA isolated from male nectaries (Figure 3). The genes chosen for qRT-PCR validation primarily included those predicted to be involved in carbohydrate metabolism and/or orthologs of genes known to be involved in nectary function. A majority of the selected genes displayed upregulation at one or more specific time-point. For example, *Starch Branching Enzyme 2 (SBE2),* encoding a protein predicted to be involved in starch synthesis, displayed high expression at the -24 hr time-point, and declined at each subsequent developmental time-point. Conversely, a gene associated with starch breakdown, *β-amylase 1 (BAM1),* displayed low expression level at every developmental stage except for the 0 hr time-point. Presumptive orthologs of several genes known to be required for nectar production in other species – *MYB305, SWEET9,* and *CWINV4* (Liu et al., 2009; Ruhlmann et al., 2010; Liu and Thornburg, 2012; Lin et al., 2014; Wang et al., 2014; Schmitt et al., 2018) – also displayed stage-specific induction. Both the *MYB305* and *SWEET9* transcripts displayed low expression at the -24 hr time-point and peaked at the 0 hr time-point, whereas *CWINV4* displayed highest expression at the -15 hr time-point.

**Figure 3.**
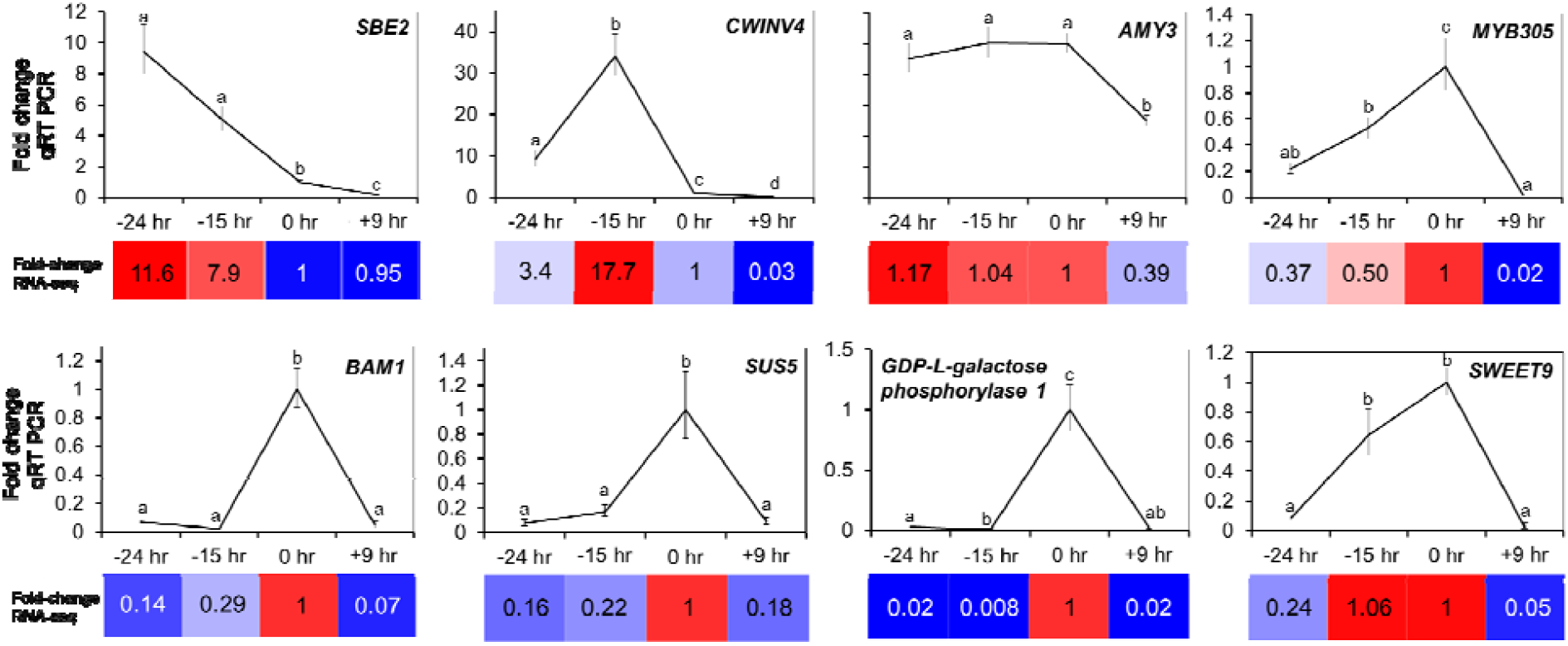
Validation of RNA-seq data. qRT PCR was used to validate the expression patterns of select differentially expressed genes in male nectaries identified through RNA-seq analyses. The fold-change in each stage was determined relative to 0 hr [the secretory stage (anthesis) was set as 1]. Statistically significant differences in expression between the stages are noted by different letters (pairwise, two-tailed T-test, p<0.05). The colored boxes represent the corresponding fold-change observed by RNA-seq analysis. SBE2 = Starch Branching Enzyme 2, CWINV4 = Cell Wall Invertase 4, AMY3 = Amylase 3, BAM1 = Beta-amylase 1, SUS5 = Sucrose Synthase 5. MYB305 and SWEET9 are the full gene names.

### 3.4 Gene ontology analyses identify a shift from anabolic to catabolic processes

Genes displaying stage-enriched expression [from Figure 2A, B and D (female, male and female/male together, respectively)] were subjected to gene ontology (GO) analyses to identify processes and metabolic pathways overrepresented in nectaries during a specific stage of development (Supplemental Files 7-9). At the -24 hr time-point, female nectaries displayed significant enrichment for anabolic processes (see *Biological Processes* and *Molecular Function* tabs in Supplemental File 7), notably transcripts encoding proteins involved in amino acid biosynthesis, transcription, and translation processes are overrepresented. However, by -15 hr, a distinct shift toward catabolic processes was noted in female nectaries, particularly with regard to starch degradation. The secretory nectaries (0 hr time-point) displayed an extension of catabolic processes to amino acids. Unique GO terms enriched at the post-secretory female nectaries (+9 hr time-point) included the appearance of those related to senescence, including assembly of the autophagosome.

Interestingly, male nectaries displayed no significant enrichment of genes related to specific GO terms at either the -24 or -15 hr time-points, which could be due to the relatively few number of genes being upregulated specifically at either stage as compared to female nectaries (Figure 2). As such, we analyzed genes commonly upregulated at the -24 and -15 hr time-points over 0 and +9 hr (i.e. the 569 genes represented in the overlapping region of -24 and -15 hr in Figure 2B, Supplemental File 8). This analysis identified both starch biosynthetic and catabolic processes as being highly enriched in pre-nectar secretion stages (i.e., -24 and -15 hr time-points). Enriched GO terms at the secretory (0 hr) and post-secretory (+9 hr) time-points of male flowers were similar to those identified in female nectaries. The GO analysis of combined reads for both male and female nectaries (from Figure 2D) closely mirrored such analyses of the male nectaries (Supplemental File 9).

### 3.5 Carbohydrate and nectar secretion-related processes

The role of starch and sugar metabolism is well characterized in nectar production (Nepi et al., 1996; Nepi et al., 2001; Ren et al., 2007; Ren et al., 2007), and this is clearly reflected in the GO analyses of the nectary transcriptomics data. The RNAseq data were characterized to identify regulation of carbohydrate interconversions as indicated by stage-specific expression of candidate genes involved in canonical biosynthetic or catabolic processes (Figure 4, Supplemental File 10). The expression of genes involved in starch synthesis primarily peaked at the -24 hr time-point (e.g. *CpSBE2* in Figures 3 and 4). There was a stark transition toward starch degradation between the -15 and 0 hr time-points, with most degradative genes having highest expression at -15 hr, the notable exception being *CpBAM1*, which was strongly induced at the 0 hr time-point (Figures 3 and 4B).

**Figure 4.**
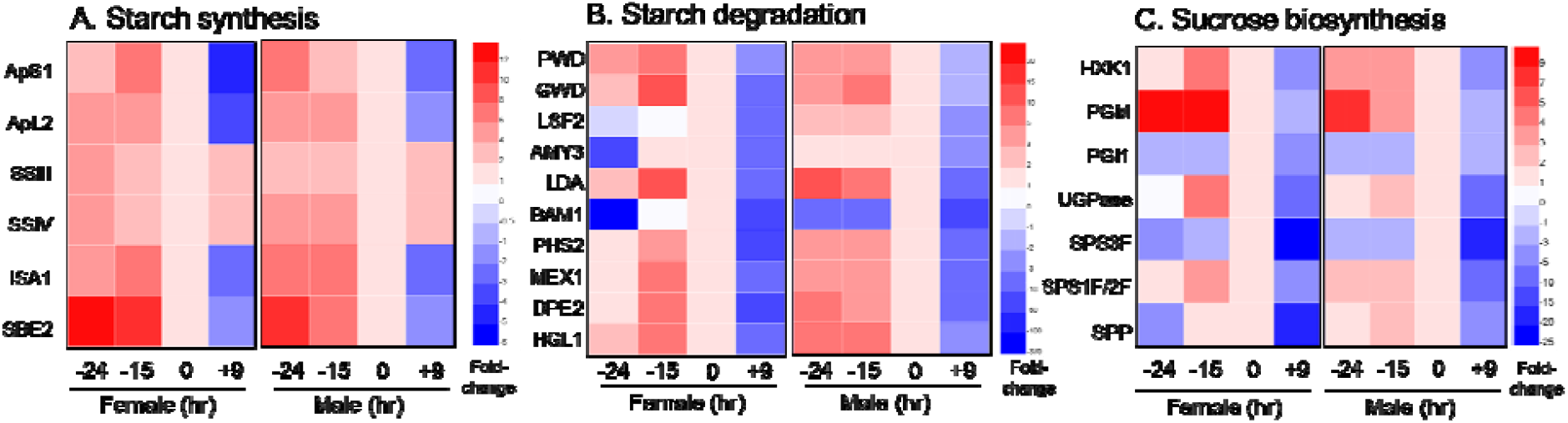
Expression analysis of genes involved in starch and sugar metabolism. Normalized RNA-seq data was used to generate heat maps for nectary-expressed genes involved in starch synthesis (A), starch degradation (B), and sucrose biosynthesis (C). The fold-change in each stage was determined relative to 0 hr [the secretory stage (anthesis) was set as 1]. In general, starch biosynthesis genes peak at -24 hr, starch degradation genes peak at -15 hr (with the exception of BAM1), and sucrose biosynthesis genes are highest at -15 and 0 hr. This analysis supports a progression of starch synthesis, starch degradation and sucrose biosynthesis as being essential for nectary function. Full names for the abbreviations of individual genes are provided in Supplemental File 10.

Sucrose synthesis coincides with starch degradation in the current model of nectar secretion (Lin et al., 2014; Roy et al., 2017). Indeed, transcript for the sucrose biosynthetic gene *Sucrose-phosphate synthase 2 (CpSPS2)* in female nectaries was relatively low at -24 hr, with high levels being observed at both -15 and 0 hr; however, in male nectaries *CpSPS2* expression was relatively constant at the -24, -15 and 0 hr time-points in male nectaries, with a sharp drop-off post-secretion (+9 hr) (Figure 4C). Expression of other sucrose biosynthetic genes generally peaked at the -15 and 0 hr time-points (Figure 4C).

Not represented in the carbohydrate metabolic pathways presented in Figure 4 are genes involved in establishing and maintaining sink status. Raffinose-family oligosaccharides (RFOs) are the primary transport sugars in cucurbits, which are hydrolyzed in sink tissues by α-galactosidases, releasing galactose and sucrose for further metabolism. Indeed, two genes encoding putative α-galactosidases, *Raffinose Synthase 6 (CpRAFS6)* and *Seed Imbibition 2 (CpSIP2)* are highly expressed in nectaries at -24 and -15 hr (Supplemental Tables 1-3), suggesting they are important for maintaining sink status during the starch filling stages.

The last two steps of nectar secretion in the current model are sucrose export via SWEET9 and extracellular hydrolysis by cell wall invertases. For the sucrose export step, *CpSWEET9* transcript began low at -24 hr and peaked at the 0 hr time-point in both male and female nectaries (Figure 3). However, the most highly expressed cell wall invertase in squash nectaries, *CpCWINV4 (Cell Wall Invertase 4),* the apparent ortholog to Arabidopsis *AtCWINV4* required for nectar secretion (Ruhlmann et al., 2010), displayed a unique timing of expression relative to other species. Specifically, its expression peaked at -15 hr (instead of at anthesis like other species), with almost no detectable transcript at either the -24 or 0 hr time-points. Lastly, with regard to regulation of nectar secretion, *CpMYB305* is the ortholog of a transcription factor intimately tied to regulating carbohydrate metabolism in *Nicotiana* and Arabidopsis nectaries (Liu et al., 2009; Liu and Thornburg, 2012; Wang et al., 2014; Schmitt et al., 2018); it displayed highest expression at the 0 hr time-point in both female and male nectaries (Figures 3).

The biological relevance of carbohydrate metabolism-related gene expression patterns was partially validated by examining starch and soluble sugar accumulation patterns and conducting enzymatic assays (Figure 5). The -24 hr and -15 hr nectary stages displayed intense iodine staining (indicative of starch) and averaged ~10-15% wt/wt starch on a fresh weight basis (Figure 5A and B). This starch was largely absent in secretory nectaries (0 hr) and remained low at +9 hr (Figure 5A and B). Consistent with a degradative mechanism, the observed decrease in starch content coincided both with an induction of β-amylase activity (Figure 5C) and the higher accumulation of soluble sugars (sucrose and glucose) in the nectaries at the 0 hr time-point (Figure 5D).

**Figure 5.**
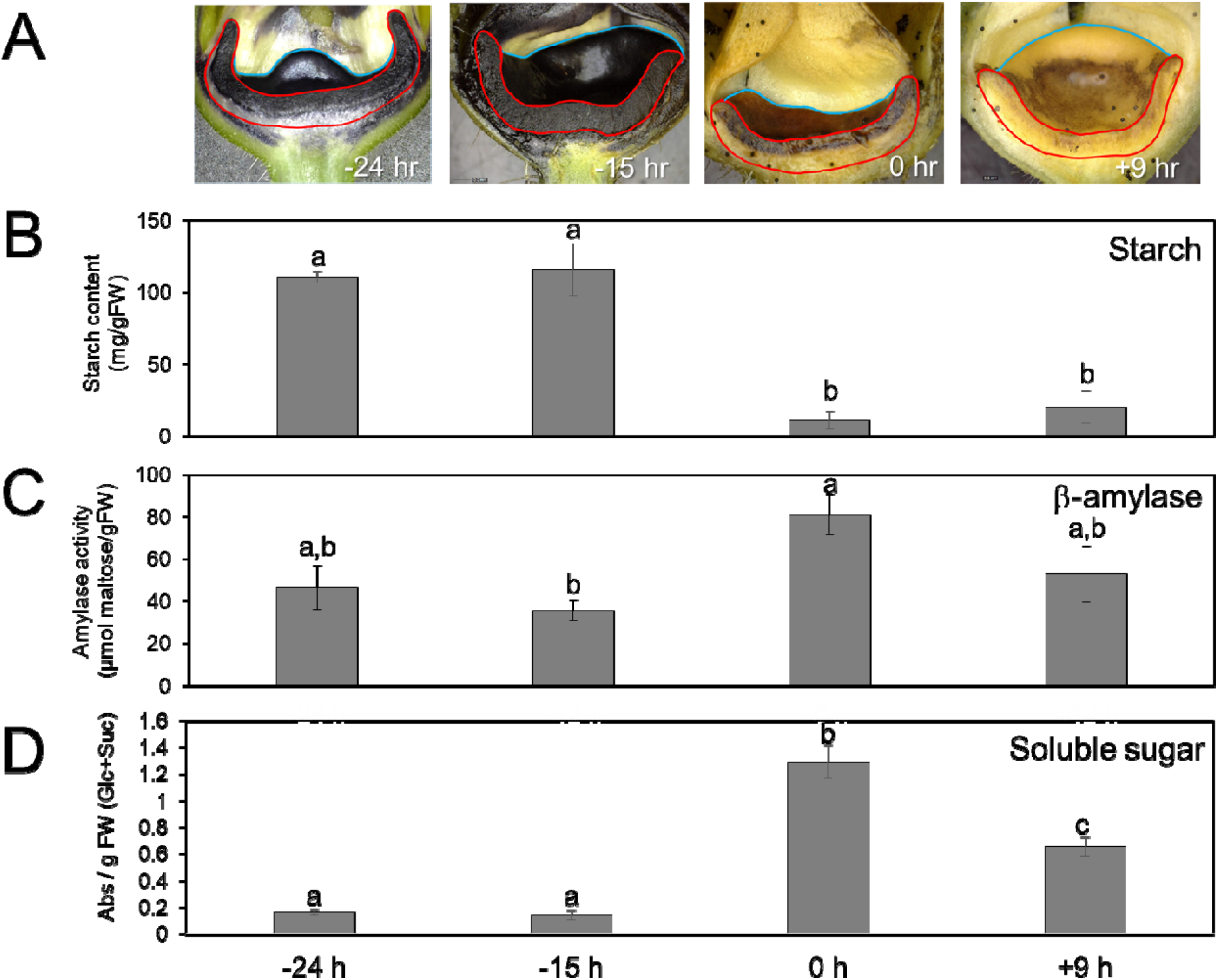
Starch and sugar metabolism in male squash nectaries at different developmental stages. (A) Longitudinally-sectioned male squash flowers stained for starch accumulation with Lugol’s Iodine solution at different developmental stages. The face of the cut nectary surface is outlined in red, whereas the remaining intact bowl-shaped nectary is outlined in blue. (B) Quantitative determination of starch in squash nectaries. (C) β-amylase activity present in squash nectaries at each stage. (D) Total soluble sucrose and glucose present in nectary tissues at each stage. Statistically significant differences between the stages are noted by different letters (pairwise, two-tailed T-test, p<0.05).

### 3.6 Transcriptional differences between female and male nectaries

Male and female flowers display a slightly different timing of nectar secretion and amount of nectar made (female flowers generally open a little later than male flowers and also produce more nectar (Nepi et al., 2001). As such, DESeq analysis was used to identify differentially expressed genes between female and male flowers specifically at the 0 hr (secretory) time-point (Supplemental File 11). This analysis identified 134 genes more highly expressed in female over male nectaries at the 0 hr time-point. In particular, the top three genes more highly expressed in female nectaries were all transcription factors, including *SHI-related sequence 1, heat stress transcription factor A-6b* and *UNUSUAL FLORAL ORGANS-like.* Conversely, male nectaries at the 0 hr time-point expressed 43 genes more highly than their female counterpart.

### 3.7 Nectar metabolite analyses

Nectars not only contains sugars, but also have many solutes with biologically important roles (Irwin et al., 2004; Nicolson and Thornburg, 2007; Elliott et al., 2008; Irwin and Adler, 2008; Adler and Irwin, 2012; Richardson et al., 2015). Nectar from both female and male flowers were characterized via a non-targeted metabolomic strategy (results shown in Supplemental Table 2). Not surprisingly, sucrose (~2.1M), glucose (~0.3M), and fructose (~0.3M) were the primary metabolites observed in both male and female nectars, with a sucrose-to-hexose ratio of ~4:1 in female nectar and ~3.4:1 in male nectar. Lesser levels of other sugars and sugar alcohols were also observable, including inositol, maltose, galactose, arabinose, erythritol, erythrofuranose, ribose, xylose, xylitol, seduheptalose and mannose (Supplemental Table 2). Additional metabolites identified in both male and female nectars included amino acids and other polar compounds, such as 2,3-butanediol, ethylene glycol, and the neurotransmitters gamma-hydroxybutyric acid (GHB) and gamma-amino butyric acid. Not surprising for an aqueous biological medium, non-polar metabolites were fewer and of lower concentrations, these included 4-coumaryl alcohol; 4-hydroxybenzyl alcohol (gastrodigenin); 2-thiophenecarboxylic acid, 4-methoxyphenyl ester; and 4-(hydroxymethyl)-2-methoxyphenol.

A handful of these metabolites differentially accumulate between male and female nectars (Figure 6, Supplemental Table 2), with most preferentially accumulating in the male nectar (~2-4-fold higher levels), including: galactose, ethylene glycol, glyceraldehyde, erythrofuranose, maltose, erythritol and tyrosol. While not statistically significant, one non-proteinaceous amino acid, gamma-amino butyric acid (GABA), occurs at 5-fold higher level in female nectar over male nectar (Supplemental Figure 1). The transcript of a putative GABA exporter (ortholog of *C. melo* XM_008448565.1) displayed a parallel increased level of accumulation in female nectaries at the secretory stage (0 hr time-point) (Supplemental Figure 1), which could account for the observed differences in GABA levels between male and female nectar.

**Figure 6.**
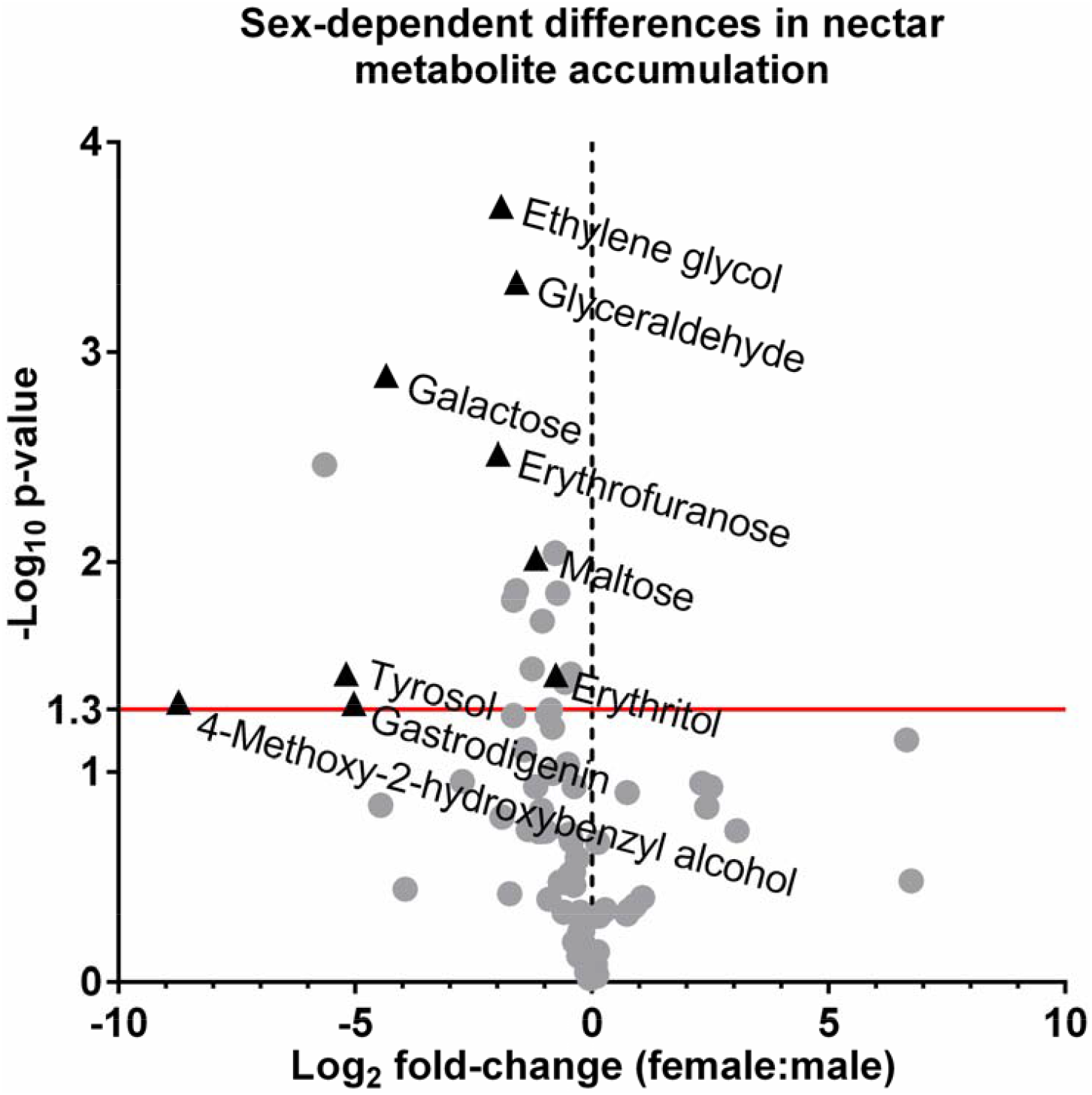
Volcano plot of the *Cucurbita pepo* nectar metabolome with the x axis representing the log2 fold change of male-to-female metabolite concentration and the y-axis representing the negative log_10_ of the adjusted p-value. Points above the red line represent metabolites with p-values < 0.05 between male and female metabolite.

## 4 DISCUSSION

This study represents the first report of global gene expression profile of cucurbit nectaries. While Arabidopsis and *Nicotiana* spp. have been extensively studied as genetic models for nectar production (Carter et al., 1999; Carter and Thornburg, 2000, 2003; Thornburg et al., 2003; Carter and Thornburg, 2004; Carter and Thornburg, 2004, 2004; Naqvi et al., 2005; Carter et al., 2006; Carter et al., 2007; Horner et al., 2007; Ren et al., 2007; Ren et al., 2007; Kram and Carter, 2009; Kram et al., 2009; Liu et al., 2009; Hampton et al., 2010; Ruhlmann et al., 2010; Bender et al., 2012; Liu and Thornburg, 2012; Bender et al., 2013; Lin et al., 2014; Stitz et al., 2014; Wiesen et al., 2016; Roy et al., 2017; Thomas et al., 2017), an expansion of molecular biology approaches into other systems with larger nectaries (Figure 1) that produce copious amounts of nectar will aid our understanding of nectary biology, particularly with regard to quantitative biochemical, physiological and comparative studies. Our study has revealed a plethora of squash genes and metabolic processes that are temporally regulated as the nectary progresses from pre-secretion to secretion to post-secretion stages of development.

Squash nectary RNAseq reads mapped to 8,863 unique *C. melo* genes, which is consistent with the number of genes identified as being expressed in Arabidopsis nectaries (9,066) and pennycress (12,335) nectaries (Kram et al., 2009; Thomas et al., 2017). Many genes displayed preferential expression at one or more stages of nectary maturation (Figure 2), with the data being validated by qRT-PCR analysis of eight targeted genes (Figure 3); demonstrating the quality and utility of the RNAseq dataset for downstream analyses.

Analysis of our transcriptome data readily identified representative candidate genes displaying the appropriate timing of expression and/or activities associated with each of the proposed steps of nectar synthesis described in previous sections, including starch synthesis, starch degradation, sucrose synthesis and sucrose export (Figures 3–5, with a summarized model in Figure 7). For example, expression of the starch biosynthesis gene *Starch Branching Enzyme2 (CpSBE2)* was highest at -24 hr and steadily decreased thereafter, which was inversely proportional to *BAM1* expression (Figures 3, 4 and 7B). In terms of maintaining sink status, there are many proteins involved in ensuring a continuous supply of photosynthate to sink tissues. There have been a few studies on carbon partitioning and resource allocation in developing cucurbit fruits (Schapendonk and Brouwer, 1984), however, little is known about the regulation of sink-source relations in *C. pepo* nectaries. Cucurbits, such as squash and cucumber, transport photosynthate in the form of raffinose-family oligosaccharides (RFOs) loaded and unloaded into the phloem predominantly via a symplastic route. In our study, we identified two very highly expressed α-galactosidases *(CpRAFS6* and *CpSIP2*) that are likely involved in establishing and maintaining sink status in nectaries.

**Figure 7.**
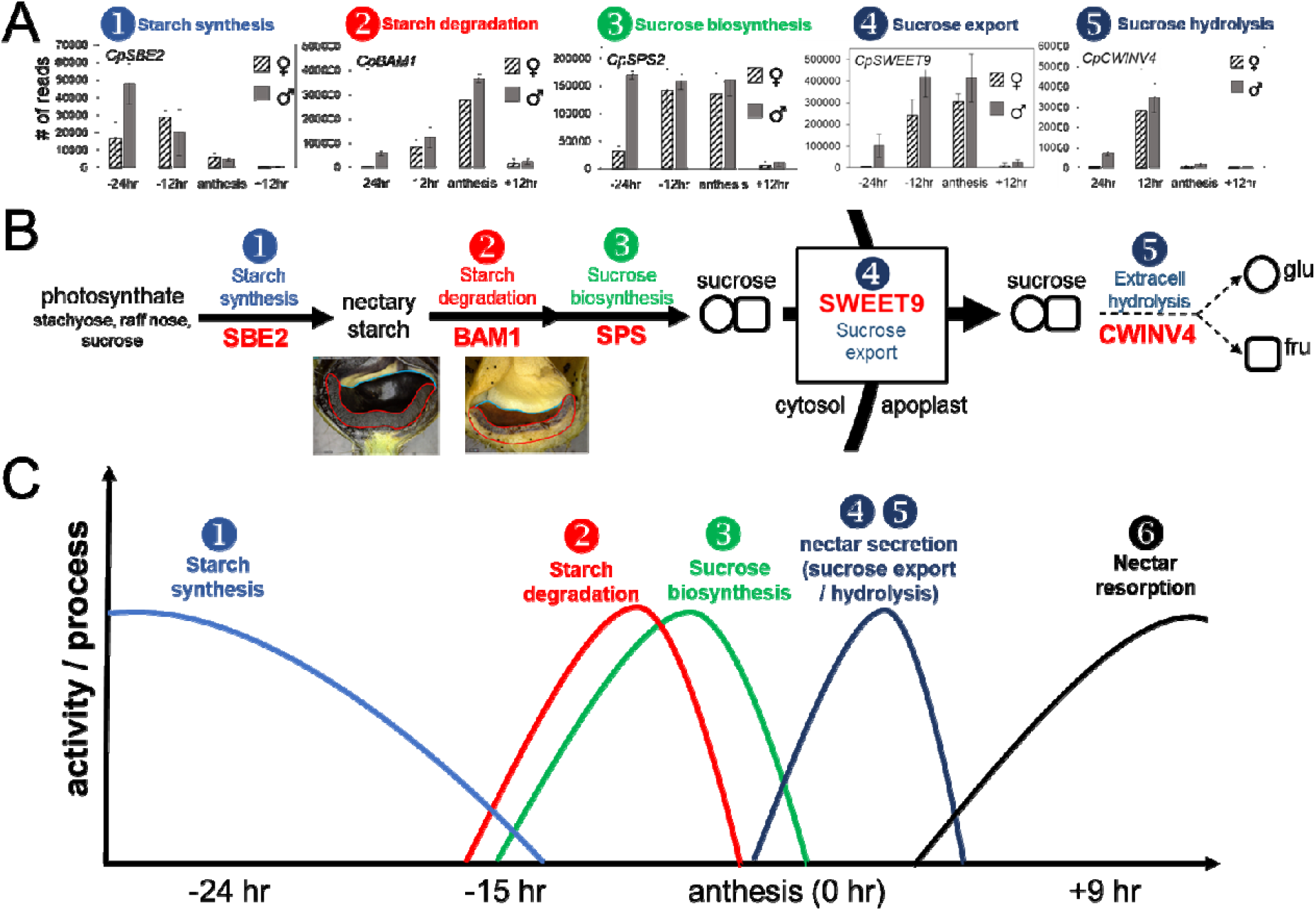
Representative genes and processes putatively involved in carbon flux in squash nectaries. **(A)** Expression of genes in male and female nectaries involved in each of the proposed steps of nectar synthesis and secretion as outlined in panel **B**; SBE = starch branching enzyme; BAM1 = b-amylase; SPS = sucrose-phosphate synthase; SWEET9 = sucrose transporter; CWINV4 = cell wall invertase **(C)** Approximate timing of processes involved in nectar production based on data from this study and previously published reports.

While the expression profiles for each of the genes and steps for nectar secretion in this study are largely consistent with what has been observed in other species [reviewed in (Roy et al., 2017)], a notable exception was *CpCWINV4.* In both Arabidopsis and pennycress nectaries, *SWEET9* and *CWINV4* have nearly identical read counts (Kram et al., 2009; Ruhlmann et al., 2010; Bender et al., 2013; Lin et al., 2014; Thomas et al., 2017); however, in squash, *CpCWINV4* is lowly expressed (maximum of ~30,000 reads) in nectaries relative to *CpSWEET9* (~400,000 reads) (Figure 7A). In contrast to Arabidopsis, *C. pepo* flowers produce nectar rich in sucrose (Nepi et al., 2001), which necessitates a reduction in extracellular sucrose hydrolysis after sucrose export from the cell. Thus, *CpCWINV4* probably does not play the same role in squash as *AtCWINV4* does in Arabidopsis (i.e. in generating a concentration gradient to drive sugar export); however, since squash nectar contains ~20% hexoses, its activity likely is important in dictating final nectar quality. It should also be noted that in sunflower a reduction in nectary CWINV expression led to an increase in nectar sucrose (Prasifka et al., 2018).

Cumulatively, it appears that modulation of cell wall invertase expression or activity represents a key step in regulating final nectar quality across species. If indeed sucrose transport across the plasma membrane is fully concentration-dependent (passive transport via SWEET9), and the final nectar sucrose concentration is >2M in squash (Supplemental Table 2), then cytosolic sucrose in the nectary parenchyma must reach at least the concentration present in the nectar. This secretory process would present a tremendous osmotic stress to the nectary tissue, which may be why senescence-related transcripts are upregulated in post-secretory (+9 hr) nectaries (Supplemental Files 7-9). A model of the approximate timing for the transition from starch synthesis to starch degradation to sucrose synthesis and export are shown in Figure 7C; since post-secretory hydrolysis likely plays a limited role in the secretory process, CpCWINV4 activity is denoted with a light dashed line.

As previously noted, nectars are much more complex than simple sugar-water, containing many classes of biologically relevant solutes that influence not only pollinator visitation, but also impact microbial growth (Nicolson and Thornburg, 2007; Roy et al., 2017). In this study, we reliably identified over 40 metabolites in both male and female nectar, which is similar to an analysis of *C. maxima* (pumpkin) nectar (Chatt et al., 2018). While male nectar preferentially accumulates certain metabolites as compared to female nectar (Figure 6), the causes or consequences of these differences are not clear. However, there were some notable findings within the metabolite analysis, including the fact that female nectar contained ~5-fold more GABA than male nectar, which was correlated to the expression of a putative GABA transporter (Supplemental Figure 1). GABA is a neurotransmitter previously reported to accumulate in squash nectar (Nepi et al., 2012), but its role in plant-pollinator interactions is currently unclear.

Along these lines, we identified another previously unreported neurotransmitter, gamma-hydroxybutyric acid, in both male and female nectars, the role of which in nectar is also unknown. Of notable interest will be to determine factors that lead to differential transcription between female and male nectaries that may lead to differences in nectar quality. In our study *MYB305,* a key transcription factor in eudicot floral nectary function (Liu et al., 2009; Liu and Thornburg, 2012; Wang et al., 2014; Schmitt et al., 2018), displayed little difference in expression between squash male and female nectaries; however, three transcription factors – *SHI-related sequence 1, heat stress transcription factor A-6b* and *UNUSUAL FLORAL ORGANS-like* – displayed preferential expression female over male nectaries. Interestingly, *SHI-related sequence 1* is a member of the *STYLISH* gene family, which control nectary development in Aquilegia (Min et al., 2018).

To summarize, our analyses have revealed a distinct profile of starch biosynthetic and degradative pathways as the nectaries progress through maturation. Our expression data reveal a similar increase in sucrose synthesis in nectaries at anthesis, as well as a number of Arabidopsis gene homologs known to impact nectary function. We provide a physiological basis behind our gene expression data with starch, amylase, and soluble sugar assays showing degradation of starch correlated with increased amylase activity and accumulation of soluble sugar. Cumulatively, our data supports the existing model of nectar secretion in the eudicots (Lin et al., 2014; Roy et al., 2017). Lastly, we have identified novel nectar metabolites that should be evaluated for their roles in plant-biotic interactions. This study represents an important step towards improving our understanding of nectar production and secretion in the cucurbits.

## AUTHOR CONTRIBUTIONS

All authors conceived and planned the study. PMK determined nectary stages and conducted RNA isolations. MH processed all RNAseq data and conducted informatics analyses in conjunction with ES, RR and CJC. ES and RR conducted starch and sugar analyses. ES conducted the qRT-PCR analyses. EC and BJN conducted the metabolomic analyses. ES, RR and CJC wrote the manuscript with feedback from all the coauthors.

## Supporting information

## ACKNOWLEDGMENTS

This work was funded by a U.S. National Foundation grant, 1339246, to MH, BJN and CJC.

---

**Supplemental Figure 1**. Female squash nectaries secrete higher amounts of GABA (A) and express higher levels of a putative GABA transporter (B) than male nectaries.

**Supplemental Table 1** – Oligonucleotides used in this study.

**Supplemental Table 2** – Untargeted total metabolite analysis of *C. pepo* nectar

**Supplemental File 1** – Read counts for all RNAseq data mapped to *C. melo* and Arabidopsis

**Supplemental File 2** – summarized read counts and DESeq analysis of nectaries

**Supplemental File 3** – List of differentially expressed genes by stage in female nectaries (by Arabidopsis hit for GO analyses)

**Supplemental File 4** – List of differentially expressed genes by stage in male nectaries (by Arabidopsis hit for GO analyses)

**Supplemental File 5** – List of commonly differentially expressed genes by stage in female and male nectaries (by Arabidopsis hit for GO analyses)

**Supplemental File 6** – List of differentially expressed genes by stage in male and female nectaries when summed together by gene (by Arabidopsis hit for GO analyses)

**Supplemental File 7** – GO analysis of differentially expressed genes in female nectaries.

**Supplemental File 8** – GO analysis of differentially expressed genes in male nectaries.

**Supplemental File 9** – GO analysis of commonly differentially expressed genes in both male and female nectaries.

**Supplemental File 10** – Carbohydrate metabolism genes and associated expression used to make heat map for Figure 4.

**Supplemental File 11** – Differentially expressed genes between female and male nectaries at the secretory stage (0 hr).

